# GATTACA: Lightweight Metagenomic Binning with Compact Indexing of Kmer Counts and MinHash-based Panel Selection

**DOI:** 10.1101/130997

**Authors:** Victoria Popic, Volodymyr Kuleshov, Michael Snyder, Serafim Batzoglou

## Abstract

We introduce GATTACA, a framework for rapid and accurate binning of metagenomic contigs from a single or multiple metagenomic samples into clusters associated with individual species. The clusters are computed using co-abundance profiles within a set of reference metagnomes; unlike previous methods, GATTACA estimates these profiles from k-mer counts stored in a highly compact index. On multiple synthetic and real benchmark datasets, GATTACA produces clusters that correspond to distinct bacterial species with an accuracy that matches earlier methods, while being up to 20*×* faster when the reference panel index can be computed offline and 6*×* faster for online co-abundance estimation. Leveraging the MinHash technique to quickly compare metagenomic samples, GATTACA also provides an efficient way to identify publicly-available metagenomic data that can be incorporated into the set of reference metagenomes to further improve binning accuracy. Thus, enabling easy indexing and reuse of publicly-available metagenomic datasets, GATTACA makes accurate metagenomic analyses accessible to a much wider range of researchers.

## 1 Introduction

Despite their important role, microbes constitute the dark matter of the biological universe. Thousands of species live in the human gut, but only a small fraction can be isolated and studied in a laboratory and very little is known about those that cannot be cultured. The short read lengths of modern sequencing instruments – combined with various inherent difficulties associated with complex bacterial environments– make it very difficult to perform simple tasks such as accurately identifying bacterial strains, recovering their genomic sequences, and assessing their abundance. Many approaches have been proposed to address these shortcomings. Specialized library preparation techniques such as Hi-C or synthetic long reads are often very accurate, but also prohibitively complex. As a result, approaches based on contig binning are more popular in practice. Metagenomic binning refers to the problem of grouping together partially assembled sequence fragments (or contigs) that belong to the same species. Current binning techniques fall into mainly two categories: (1) supervised classification of contigs into known taxons via comparisons to previously catalogued species [13, 26, 30] and (2) unsupervised clustering techniques using features derived directly from the metagenomic sample data [**?**, 2, 3, 16, 17, 20, 31], where unsupervised clustering has the clear advantage of binning contigs that pertain to previously unknown species. While some unsupervised techniques [17, 28] perform clustering based only on the contig sequence composition (the frequency of certain short motifs, e.g. all tetra-mers), the most successful recent approaches [2, 3, 16, 20, 31] also incorporate contig coverage profiles across multiple metagenomic samples. In brief, these techniques assemble de-novo bacterial contigs and estimate the coverage of each contig within each sample of a large mategenomic cohort using read mapping. Naturally, contigs belonging to the same species will have similar abundances across different samples (determined by which cohort samples the species is present in); coverage profiles can therefore be used to cluster related contigs. This approach is accurate but has two main limitations: it requires a large cohort of samples, as well as sizable compute resources for read alignment. We address both of these limitations in this work.

In particular, we present GATTACA, a lightweight framework for metagenomic binning, which (1) avoids read alignment without loss of accuracy and (2) enables efficient stand-alone analysis of single metagenomic samples. Both results are based on the finding that we can approximate contig coverages using kmer counts while still achieving the same binning accuracy as leading alignment-based methods. In addition to offering a significant speedup in coverage estimation, using kmer counts, as opposed to alignment, provides us with the exciting ability to index *offline* any publicly-available metagenomic sample and incorporate it into the coverage profile of the contigs being processed. This allows us to efficiently pull in data from large growing repositories, such as the Human Microbiome Project (HMP) [29] or EBI Metagenomics archive [12] into any metagenomic study (especially one where only a single or few samples are available) at almost no cost. For example, our kmer count index for a typical HMP sample only requires 100MB on average. We achieve the small space requirement by leveraging memory-efficient hashing with minimal perfect hash functions (MPHFs) and the probabilistic Bloom filter data structure. In contrast, using these datasets with read alignment would require massive downloads (for example, a single HMP sample is roughly 7GB compressed and 30GB uncompressed) and expensive subsequent handling to map the reads. In terms of speedup, we found our coverage estimation time to be at least an order of magnitude faster (approximately 20*×*) when the index is computed offline (e.g. for recyclable public reference samples) and about about 6*×* when the kmers are counted on-the-fly (e.g. for private samples used only once), when compared to read mapping.

While using small indices allows us to incorporate a large number of publicly-available samples into a given study, not all existing samples will carry content relevant to the study in question. Namely, samples that don’t contain any of the species present in a given set of contigs cannot contribute any useful information for grouping the contigs. The same logic applies also to samples that carry content identical to a sample that has already been included. This motivates the need to additionaly define appropriate sample selection criteria, for which we propose two metrics: (1) relevance and (2) diversity. More specifically, we would like to select a panel of samples which share content with the sample being analyzed (our query) but that also differ in the content that is shared. We use locality sensitive hashing [15] and the MinHash technique [7], to compare the samples efficiently. At a high level, we create and index small MinHash fingerprints for each sample in the database (offline), and then extract the appropriate samples according to the fingerprint of the query. The resulting index can be separately downloaded and used to determine which samples to include into the analysis; it needs to be updated only occasionally when new samples become available.

We evaluate GATTACA in clustering contigs assembled across multiple samples (co-assemblies) and from individual samples, using both synthetic and real datasets. We compare our results with several state-of-the-art methods in metagenomic binning: CONCOCT [3], MetaBat [16], and MaxBin [31], using standardized cluster evaluation metrics and benchmarks (reusing evaluation scripts from existing methods when appropriate). GATTACA was implemented in C++ and Python and is freely available at http:://viq854.github.com/gattaca.

## 2 Methods

### 2.1 Index of Kmer Counts

In order to quickly estimate contig coverages, GATTACA builds a small index of kmer counts for each sample in the cohort. Several solutions have been proposed for exact (e.g. using hash maps [21] or minimum perfect hash functions [24]) and approximate kmer counting (e.g. using the count-min sketch [32]). Since the content of each sample in our panel is static, our index uses a minimal perfect hash function [9] to store the kmer counts without loss of accuracy, resulting in a drastic reduction in space when compared to traditional hash tables (we also found it to be more space-efficient than the count-min sketch solution for the same binning accuracy). At a high level, given a set *S* of *n* keys, a minimal perfect hash function (MPHF) *h* provides a mapping between the keys and *n* consecutive integers from 0 to *n* − 1; that is, *h* is an injection on *S*, guaranteeing no collisions among its keys (for *x* and *y* in *S*, if *x* ≠*y*, then *h*(*x*) ≠ *h*(*y*)) and exactly *n* possible outputs from the integer set {0, 1, 2, *…, n-*1}. We use the BDZ algorithm based on random *r*-partite hypergraphs [6] for constructing the MPHFs.

#### Index Construction

To construct the index, we first generate the kmers from the all reads in the sample (accounting for both forward and reverse complement strands) and exclude kmers that occur only once, since these are most likely present due to sequencing errors. We use a kmer length of 31-bp in our experiments (compacting the kmers into 64-bit integers for convenience). We then generate the MPHF, *h*_*S*_, for the resulting set of distinct kmers, *S*, and store their counts in an integer array *A* (*|A|* = *|S|*), at the indices given by *h*_*S*_; namely, *A*[*h*_*S*_ (*x*)] = *count*(*x*), for each kmer *x ∈ S*. We found 8 bits to be sufficient for storing the kmer counts (and since many counts are small, these can be compressed even further using techniques such as varint encoding). Finally, we need to store the elements in *S* to support lookups, since *h*_*S*_ (*z*) for *z* ∉ *S* will return a valid but incorrect index into *A*. One direct solution for storing *S* would be to rely on the MPHF, using a secondary array *B* and setting *B*[*h*_*S*_ (*x*)] = *x* for all kmers *x* ∈ *S*; then we could check upon lookup of a key *y*, if *B*[*h*_*S*_ (*y*)] is equal *y* and determine whether *y* was in the set. However, this solution requires storing the array *B* of *|S|* 64-bit integers, which is 4× larger than *A*, and would substantially increase the index. So instead, we store the set *S* in a Bloom filter, *BF*, which is a widely used probabilistic data structure for testing set membership that offers space-efficiency at the expense of possible false positives (no false negatives are possible). We have configured the size of *BF* based on a false positive probability of 0.05. As a result, our index for each sample consists of: (1) the MPHF, *h*_*S*_, (2) the array of counts, *A*, and (3) the Bloom filter storing the elements of *S*, *BF*. As an example, the size of the index constructed for an HMP sample containing 20 million 100-bp long reads was 108MB.

#### Coverage Estimation

Given a contig *c* and an index *I* of a cohort sample, we estimate the coverage of *c* in this sample by performing lookups in *I* for each kmer in *c* and then computing the median of the resulting counts. More specifically, we return the median of the set of counts *C* = *…*`count`(*x*)…. ∀ `kmers` *x* ∈ *C*, where,

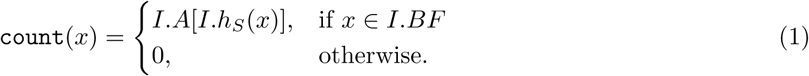

### 2.2 Contig Representation

Given a set of contigs assembled from a single or multiple metagenomic samples, our goal is to bin together the contigs that belong to the same class (e.g. species or strain). Similar to existing methods, e.g. CONCOCT, we first represent each contig as a multi-dimensional vector using both its sequence composition and coverage profile across multiple samples, where our coverages are approximated using kmer counts instead of read mapping, as described above. Namely, given *M* reference samples (either from the same study or from a public database), our coverage profile is the median count of the contig kmers in each sample., while the composition profile is the normalized frequency of each possible tetra-mer in the contig and its reverse complement (resulting in a total of *F* = 136 such features); the normalization of composition features is done according to the CONCOCT procedure (please see [3] for details). Therefore, each contig is a vector *V* = [*c*_1_,…, *c*_*M*_, *f*_1_,…, *f*_*F*_], where *c*_*i*_: = the median kmer count in sample *i* and *f*_*j*_ := the frequency of tetra-mer *j* in the contig sequence.

### 2.3 Clustering algorithm

Given the resulting vector representations, we cluster the contigs using a Bayesian Gaussian mixture model (GMM) with a Dirichlet prior. In brief, we define a mixture distribution *p* of *K* Gaussian components over *n* data points *x*_*i*_∈ ℝ^*d*^ and (unobserved) assignment labels *z*_*i*_ ∈ {0, 1}*^K^* for *i* = 1,…, *n*. Our model is the product of a likelihood term

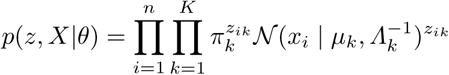

and a prior term

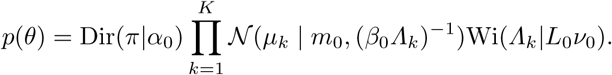

Here, *X* ∈ ℝ^*d×n*^ is the matrix of data points and 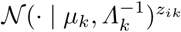 is a multivariate Gaussian with mean *μ*_*k*_ and inverse covariance matrix∧*_k_*. The *π ∈ Δ* _*K-*1_ form a vector of cluster weights. Together, the *μ*_*k*_, ∧*_k_, π* form the parameter vector *θ* of the likelihood. The prior over *θ* is a product of a Dirichlet with hyper-paremeter *α*_0_ ∈ *Δ*_*K-*1_, a multivariate normal with hyper-parameters *m*_0_ ∈ ℝ^*d*^, *β* _0_ > 0 and a Wishart distribution parametrized by *L*_0_ ∈ ℝ^*d*×*d*^ p.s.d. and *v*_0_ > 0.

We perform inference by maximizing the marginal log-likelihood log *p*(*X*) using variational inference. In brief, we maximize the evidence lower bound

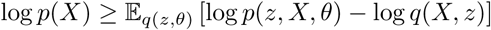

over the set of approximating distributions *q*. By our choice of conjugate prior, the posterior *p*(*θ,z|X*) and hence the optimal *q* have the same form, which factors over *q*(*z| θ*)*q*(*θ*). We optimize the bound using variational expectation-maximization, which consists of repeatedly choosing updating *q*(*z|θ*) and *q*(*θ*). Each update has a closed-form solution by our choice of conjugate prior. We conclude the algorithm by assigning each data point to its maximum *a-posteriori* label according to *q*(*z|θ*). We refer the reader to section 21.6.1 in the standard textbook of Murphy (2012) [22] for the full derivation of this algorithm.

At a high-level, the above model is very similar to automatic relevance determination (ARD), which is used by CONCOCT. We have found our approach to perform better in practice than ARD, especially for automatically determining the number of clusters in the data. Both algorithms are implemented in our software package. Other alternative clustering methods can also be easily plugged into GATTACA’s binning pipeline.

#### 2.4 Sample Selection

Given a query sample, *Q*, we would like to select *n* samples from the public database, which can provide discriminatory features for clustering the contigs of *Q* (where the features represent the coverage of the contigs in the respective samples). Intuitively, the selected samples must share some content with *Q* (have *relevance*), as well as have pairwise *diversity* among themselves to guarantee coverage of different contigs of *Q*. Similar relevance and diversity concepts can be found in online recommendation systems (e.g. for articles or music [1, 8]).

By representing each sample as a set of overlapping kmers, we apply the Jaccard coefficient to measure their similarity, where the Jaccard coefficient 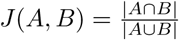, for two sets *A* and *B*. Then, we can consider relevant the samples that are within a certain distance from *Q* under Jaccard (e.g. all samples *S* for which *J* (*S, Q*) *>* 0). However, computing the Jaccard distances directly on the kmer sets would be inefficient for large databases. Therefore, we apply the *min-wise independent permutations* (MinHash) LSH scheme [7] to create small fingerprints for each sample set instead, defined as follows.

Let *U* be the ground set of all possible set items. Given a random permutation *π* of indices of *U* and a set *X*, let *h*_*π*_(*X*) = *min*_*x∈X*_ {*π*(*x*)}. The MinHash LSH family *H* will consist of all such functions for each choice of *π*. It can be easily shown that for a given *h* chosen uniformly at random, *P r*[*h*_*π*_(*A*) = *h*_*π*_(*B*)] = *J* (*A, B*) (see [7] for details). Due to the high variance in the probability of collision, we concatenate *L* different hash functions from the family *H* chosen independently at random to form the fingerprint. Then given the number of the hash collisions among the chosen *L* functions, *c*, the ratio *c/L* can also be used as an unbiased estimator for *J* (*A, B*).

To summarize, given the kmer set *K* = {*s*_0_, *s*_1_,…, *s*_*n-*1_} of some sample *S* and *L* hash functions from *H*, we construct the MinHash fingerprint vector *F* = [*f*_0_, *f*_1_,…, *f*_*L*-1_], such that the fingerprint entry *f*_*i*_ is the minimum set element under hash function *h*_*i*_:

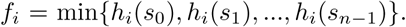

Now given the fingerprints, we can define relevance between a sample *S* and *Q* as simply the number of entries shared by their fingerprints. By indexing the fingerprints of all the samples in the database into *L* tables (based on the value of each fingerprint entry, respectively), we can find all the samples that share at least one fingerprint entry in common with *Q* using simple lookups, as well as rank them according to relevance.

Finally, if the number of relevant samples is too high, we can reduce our panel using the diversity criterion. That is, given all the relevant samples, we can select *n* samples that maximize the diversity of the set. This problem is known as the *dispersion* problem [10], where the objective is to locate *k* points among *n*, such that some function of distances between the *k* points is maximized. One popular optimality criteria is the MAX-MIN, which maximizes the minimum distance between a pair of points. This problem is known to be NP-hard; however, an efficient greedy heuristic algorithm exists for the MAX-MIN dispersion problem when the distances satisfy the triangle inequality, with provable performance guarantee of 2 [25]. Given two samples *A* and *B*, we define their diversity as: *D*(*A, B*) = 1 – *J* (*A, B*) and apply the greedy algorithm of [25] to find the *n* samples. While this procedure is simple and can be efficiently used to detect samples with distinct kmer sets, its main limitation is that it cannot be used to find samples which differ in kmer frequency only (since frequency does not affect Jaccard distance), which could also be used to generate discriminatory features.

### 3 Results

#### 3.1 Datasets

##### Synthetic datasets

We used two synthetic datasets generated by Alneberg et al. [3] from the 16S rRNA samples of the Human Microbiome Project(HMP) [29]. The first dataset (“Species-Mock”) consists of 96 samples containing a mixture of 101 different species (without strain-level variation), while the second dataset (“Strain-Mock”) consists of 64 samples comprising a mixture of 20 different organisms, of which some represent strains of the same species (e.g., this dataset includes five different *E*. *coli* strains). The relative abundance profiles of the species and strains in each sample were assigned according to the distribution of the 101 and 20 most abundant organisms in the original HMP samples, respectively. Reads (100-bp long) were simulated from random positions of the genomes present in the sample based on their relative abundance, for a total of 7.75 million reads and 11.75 million reads in each “Species-Mock” and “Strain-Mock” sample, respectively. Both datasets contain the set of contigs co-assembled across all the samples by Alneberg et al. using the Ray assembler [5], and partitioned into fragments of 10 kilobases when appropriate. We used the default minimum contig length of 1000-bp when running CONCOCT, MaxBin, and GATTACA; this parameter was set to 1500-bp for MetaBat, which is the smallest length supported by this method. As a result, the “Species-Mock” included 37,627 valid contigs and the “Strain-Mock” included 9,411 valid contigs (for all tools except MetaBat). To determine the genome assignment for each contig, the simulated reads were mapped back to the contigs and the genome was selected based on the majority of the mapped reads (with the requirement that it corresponds to at least 50% of the reads and there are at least 10 unambiguously mapped reads); this is the same criteria using by Alneberg et al.

To evaluate single-sample binning, we also assembled the contigs for each simulated sample of the “Species-Mock” independently. We used the SPADES [4] assembler with default parameters and automatic k-mer selection, and applied the same post-processing and labeling criteria.

##### Real datasets

To evaluate co-assembly based binning we used the small Sharon et al. (2013) dataset comprised of 11 fecal samples from a preterm infant [27]. We downloaded 1.372 10 × ^8^ 100-bp short reads from the SRA052203 NCBI archive as 18 separate samples (of which 7 were resequenced from the original 11 samples). The co-assembled contigs for this dataset were made public by Alneberg et al. [3]. To test binning accuracy, we used the CheckM [23] method based on single-copy core genes (SCGs), which are housekeeping genes that occur in single copies in the microbial genomes and have been previously used to evaluate the purity of the metagenomic clusters [3].

To evaluate binning of contigs assembled from single samples, we downloaded Illumina raw sequencing reads from the following two publicly-available gut microbiome sample collections: (1) all the available gut samples from HMP (148 total) and (2) 56 gut samples from the Nielsen et al. study [2] available through the European Nucleotide Archive. We used the SPADES [4] assembler with default parameters to assemble the contigs of individual samples (cutting contigs into 10 kilobase fragments and filtering contigs shorter than 1000-bp, as in simulation). Since the HMP samples were very large (over 30GB/sample uncompressed), we subsampled them using the seqtk program [11] to 20 million paired-end reads per sample during index construction. To label the contigs in each evaluated sample we used the TAXAssign program. Labels were assigned based on the top 100 hits against the BLAST NCBI genome database that had at least 95% identity and 90% query coverage. A taxonomic label was assigned if 90% of the hits at a given level belong to the taxon associated with that label. This procedure classified roughly 15% of them on average at the species level depending on the sample.

Since only a small fraction of contigs could be classified via TAXAssign, we also used SCGs to evaluate binning accuracy. In addition to CheckM, we also applied Prodigal [14] to predict and functionally annotate genes on our sample contigs and then RPS-BLAST to COG annotate the protein sequences (using the NCBI COG database). We then used the 36 Clusters of Orthologous Groups (COGs) pre-selected by Alneberg et al., along with their script COG table.py to generate the number of SCGs in each contig cluster.

#### 3.2 Clustering Evaluation Metrics

We apply the following three standardized metrics for evaluating metagenomic clustering of labelled contigs (e.g simulated contigs for which we know the ground-truth genome assignment or contigs confidently classified using TAXAssign): (1) *recall* - measures how many same-class contigs are placed in the same cluster, (2) *precision* - measures the purity of each cluster, and (3) *adjusted Rand index* (ARI) - measures how often pairs of same-class contigs are clustered together. Let *N* be the total number of contigs, *K* be the number of computed clusters, and *S* be the number of contig labels in the data (e.g. different species or strains). Also let *n _ks_* be the number of contigs in the *k*-th cluster with label *s*. Then we have:

1. 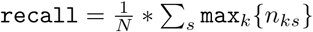
2. 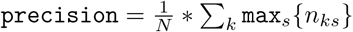
3. 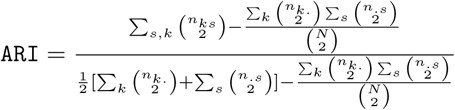 where *n*_·*s*_ = ∑_*k*_ *n*_*ks*_ and *n*_*k*·_ = ∑_*s*_ *n*_*ks*_.

### 3.3 Synthetic Benchmarks

First we show that replacing read mapping with kmer counting for coverage estimation does not result in loss of clustering accuracy using two synthetic benchmarks created by Alneberg et al. [3]. Namely, we compare the performance of GATTACA, CONCOCT, MaxBin, and MetaBat on the “Species-Mock” and “Strain-Mock” datasets, using the published co-assembled contigs. Here we build the index of kmer counts only from the cohort of samples associated with each dataset. As a result, our “Species-Mock” index contains 96 samples (with a total size of 12GB) and our “Strain-Mock” index contains 64 samples (size of 5.1 GB). Table 1 shows the accuracy results for each tool (computed with the Validate.pl script provided by CONCOCT). GATTACA achieves similar recall to CONCOCT and MaxBin (all higher than MetaBat) on both datasets, while its precision and ARI are somewhat lower on the “Species-Mock” and higher on the “Strain-Mock". MetaBat achieves the highest precision on both datasets; however, it clusters substantially fewer contigs.

**Table 1:** Binning performance on synthetic data. Synthetic datasets were obtained by generating reads from known genomes; bacteria differ either at the strain or at the species level.

| Dataset | Contigs | Labeled contigs | True clust. | Pred. clust. | Rec | Prec | Adj. Rand Idx. | Program |
| --- | --- | --- | --- | --- | --- | --- | --- | --- |
| Species-Mock | 37,628 | 37,589 | 101 | 102 | 0.993 | 0.978 | 0.971 | GATTACA |
|  |  |  |  | 101 | 0.998 | 0.988 | 0.983 | CONCOCT |
|  |  |  |  | 101 | 0.997 | 0.989 | 0.982 | MaxBin |
|  | 37,501 | 37,490 | 101 | 139 | 0.950 | 0.999 | 0.953 | MetaBat |
| Strain-Mock | 9,417 | 9,109 | 20 | 22 | 0.975 | 0.968 | 0.973 | GATTACA |
|  |  |  |  | 21 | 0.982 | 0.942 | 0.945 | CONCOCT |
|  |  |  |  | 116 | 0.967 | 0.917 | 0.879 | MaxBin |
|  | 6,281 | 6,244 | 20 | 34 | 0.856 | 0.998 | 0.857 | MetaBat |

Since one of the goals of this project is to support single-sample analysis, we also evaluate single-sample contig clustering on simulated data. In particular, we assembled several samples of the synthetic “Species-Mock” dataset separately and used the remaining samples to build the index of kmer counts in a “leave-out-out” fashion; namely, for a given sample *Q* our index includes all the samples in the original cohort except *Q*. Table 2 shows the performance on three different samples, demonstrating the accuracy of our approach.

**Table 2:** Leave-one-out accuracy of GATTACA on the “Species-Mock” dataset. We assemble contigs from a single sample and cluster them with a reference index constructed from all the other samples. Performance is comparable to when all contigs were co-assembled.

| Sample | Ctgs | Labeled Ctgs. | True clust. | Pred. clust. | Rec | Prec | Adj. Rand Idx. |
| --- | --- | --- | --- | --- | --- | --- | --- |
| S1023 | 8,245 | 8,237 | 25 | 19 | 0.964 | 0.985 | 0.952 |
| S1063 | 12,845 | 12,824 | 25 | 25 | 0.991 | 0.932 | 0.919 |
| S1095 | 12,863 | 12,848 | 31 | 28 | 0.996 | 0.995 | 0.996 |

### 3.4 Real Data Benchmarks

Next we evaluate binning on the real Sharon co-assembly. Since the true reference genomes are not available for this experiment, we use the CheckM [23] method based on SCGs to approximate recall and precision. The Sharon dataset consists of 18 samples and the resulting GATTACA index has a size of 2.2GB. Figure 1 shows the number of genomes identified by each method with a precision higher that 90% and 95%, respectively, and varying recall, as reported by CheckM. All methods, except MetaBat, produce 3 highly-accurate genomes with precision and recall ≥ 95% (MetaBat generates only one such genome). GATTACA’s performance is very similar to CONCOCT here, with the exception of an unreported bin at precision ≥ 90%, which CONCOCT produces with a recall of 30%.

**Fig. 1:**
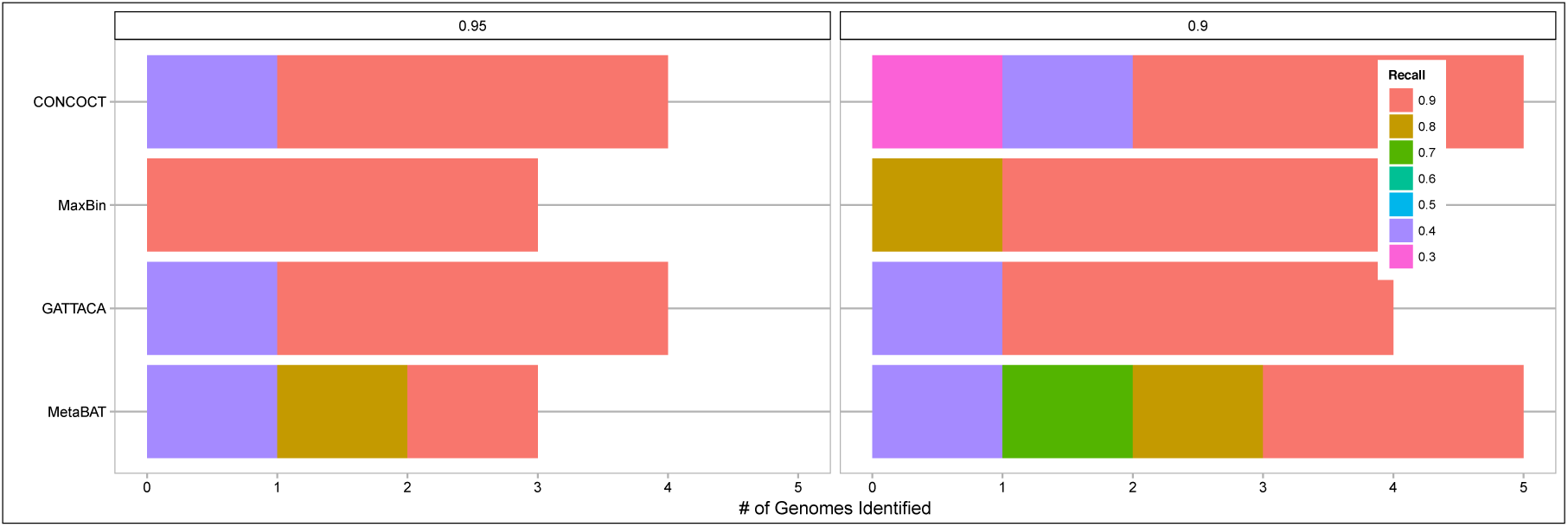
Binning performance on the real Sharon dataset reported by CheckM [23]. The number of genomes identified with ≥ 90% (left) and ≥ 95% (right) is shown for varying recall. The image was generated the ‘benchmark.R’ script (slightly modified) provided by MetaBat.

We also evaluate binning of single-sample assemblies using a panel of publicly available datasets of the gut microbiome. Table 3 shows the binning accuracy computed using TAXAssign labels for three HMP stool samples clustered using a panel of 95 different stool samples downloaded from the HMP database. Due to the long read mapping runtime, we show results for CONCOCT, MetaBat, and MaxBin on only one of the HMP samples, SRS011239. Figure 2 also shows the SCG-based results for SRS011239 obtained using CheckM.

**Fig. 2:**
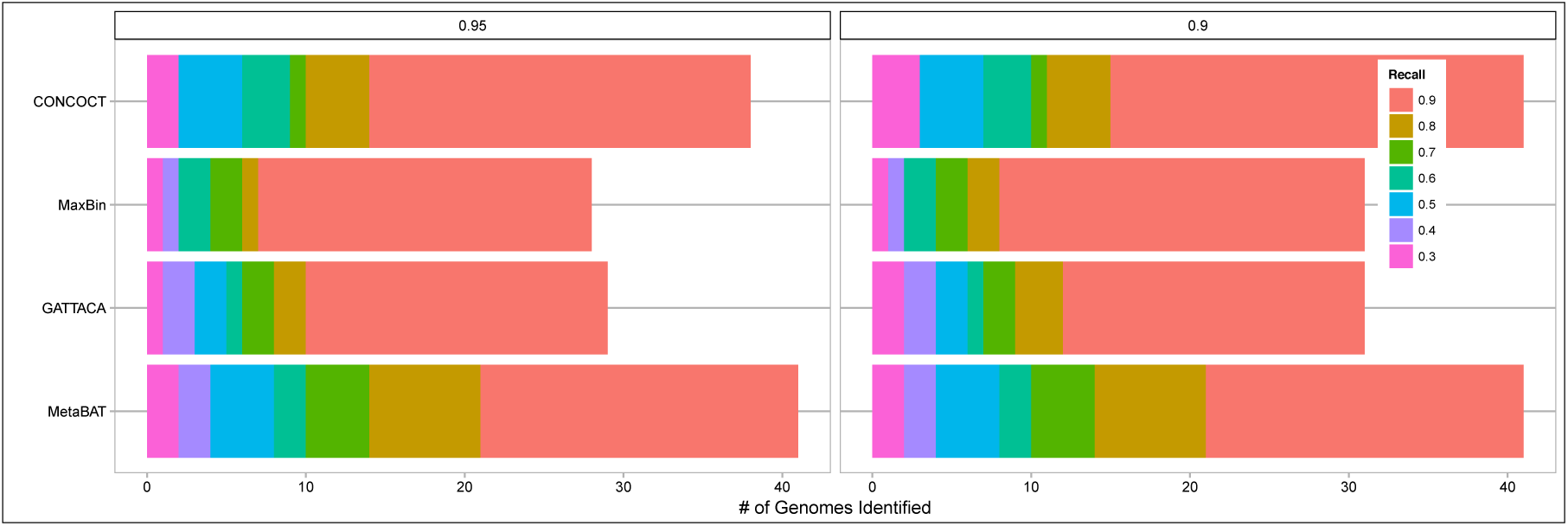
Binning performance on the real SRS011239 sample and a panel of 95 HMP metagenomes reported by CheckM [23]. The number of genomes identified with ≥ 90% (left) and ≥ 95% (right) is shown for varying recall. The image was generated the ‘benchmark.R’ script (slightly modified) provided by MetaBat.

**Table 3:** Clustering of real HMP samples using a panel of 95 other HMP metagenomes. Performance is stable and accurate across samples, with an adj. Rand index of about 90%. True assignments were obtained using TAXAassign. Contigs that were not covered by at least a single sample were not included in the clustering step.

| Dataset | Ctgs | Labeled Ctgs | True Clust. | Pred. Clust. | Rec | Prec | Adj. Rand idx. | Program |
| --- | --- | --- | --- | --- | --- | --- | --- | --- |
| SRS011239 | 28,839 | 5,430 | 46 | 131 | 0.910 | 0.963 | 0.929 | GATTACA |
|  | 33,180 | 5,456 | 48 | 212 | 0.902 | 0.959 | 0.906 | CONCOCT |
|  | 33,166 | 5,456 | 48 | 52 | 0.928 | 0.936 | 0.910 | MaxBin |
|  | 16,136 | 3,419 | 17 | 51 | 0.920 | 0.988 | 0.937 | MetaBat |
| SRS011134 | 37,645 | 5,200 | 54 | 128 | 0.848 | 0.920 | 0.892 | GATTACA |
| SRS011302 | 28,034 | 4,712 | 55 | 115 | 0.875 | 0.929 | 0.898 | GATTACA |

Next we evaluate GATTACA in clustering contigs from a single sample of the Nielsen et al. [2] study using an index built from 46 samples of the study and augmented with 83 HMP samples selected using the MinHash sample index based on the relevance criteria described above. Table 4 show the results based on the TAXAssign labels. We also show the results when using the 46 samples of the study only. It can be seen that the binning precision improves significantly with the addition of the HMP samples, motivating the inclusion of publicly-available data in studies with fewer available samples.

**Table 4:**
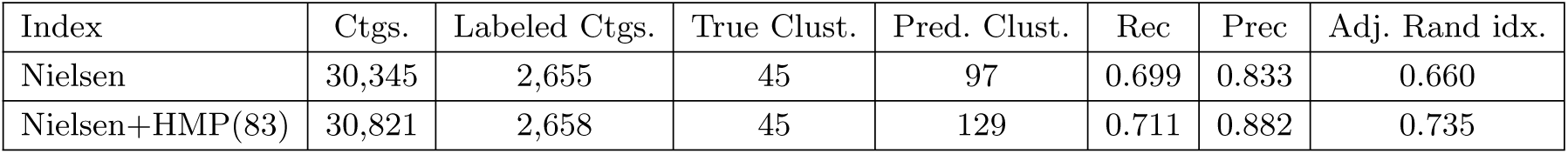
Binning sample SAMEA1965191 from the study of Nielsen et al. [2] using an index of samples from the study and an index augmented with HMP samples. Increasing the reference size greatly improves clustering accuracy. Contigs that were not covered by at least a single sample were included in the clustering step.

### 3.5 Evaluation of Sample Selection with MinHash

In order to efficiently leverage publicly-available data, we apply the MinHash technique to quickly identify public samples that are relevant (i.e. share content) to a given metagenomic study. In particular, we quickly compare the fingerprint(s) of the study’s single or multiple samples, to the fingerprints (indexed for efficiency) of publicly-available metagenomic datasets. We found that MInHash fingerprints of length 1024 provide a good tradeoff between runtime and sensitivity. This requires only 8K additional space per metagenomic sample to store its fingerprint. In particular, we performed the following experiment to evaluate the performance of MinHash for this task. Given a randomly-selected real query sample *Q* (namely, SRS011134 from the HMP), we created a panel of 50 samples (each containing 10M reads of length 100-bp) that share a varying fraction of reads with *Q*, ranging from 2% to 100%. More specifically, we first started with *Q* and 49 samples containing reads simulated from the chicken genome (under the assumption that there will be little content similarity between this genome and the species present in SRS011134). We then replaced varying fractions of the chicken reads with the reads of SRS011134. Figure 3 shows the true Jaccard similarity between the panel samples and *Q*, as well as the Jaccard similarity estimated using MinHash, with varying fingerprint lengths.

**Fig. 3:**
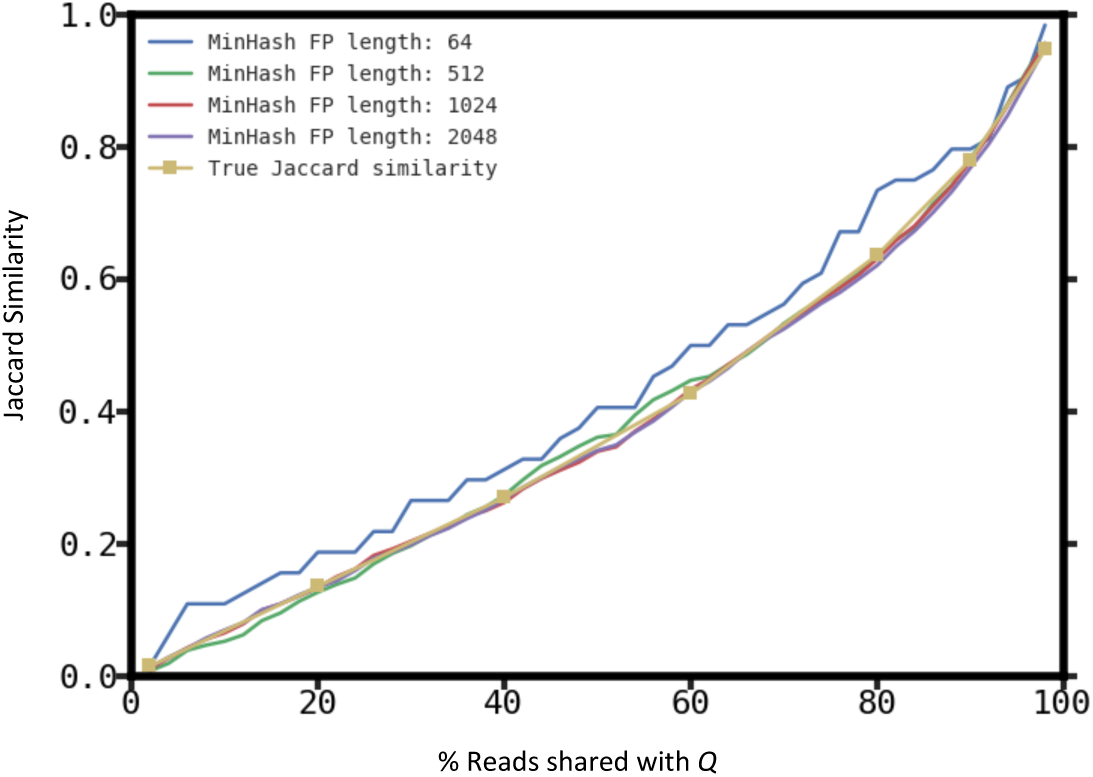
Jaccard similarities estimated using MinHash for a query sample *Q* (SRS011134) and a panel of 49 samples sharing varying levels of content similarity with *Q*. The true Jaccard similarity is shown along with estimates computed from fingerprints of different length.

To further analyze the power and efficiency of using MinHash for metagenomic sample comparison, we downloaded 32 HMP samples originating from different body sites (e.g. gut, saliva, throat). We then computed the MinHash fingerprints (of length 1024) of each sample and the Jaccard distances estimated using these fingerprints. Figure 4 shows the resulting distance matrix and hierarchical clustering of these 32 samples. As expected, samples originating from the same body site are significantly more similar in content, as reflected by the MinHash fingerprints, and cluster together. The pairwise distance computations performed in this experiment took less than a second total, and can easily scale to large datasets of publicly available metagenomic datasets; suggesting that MinHash is a good candidate for the quick and accurate metagenomic sample comparison.

**Fig. 4:**
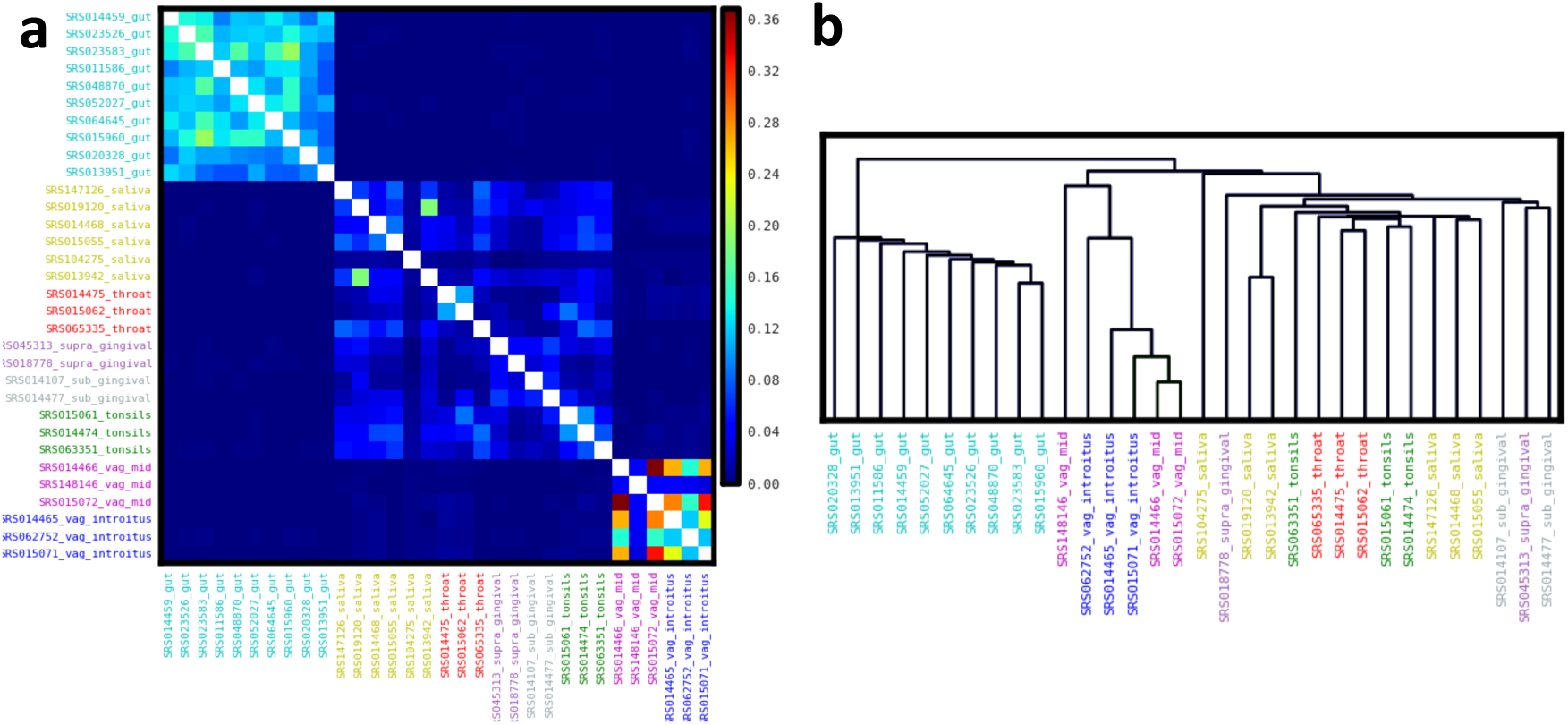
Jaccard similarities estimated using MinHash for a panel of 32 HMP samples extracted from different body sites. (a) Heatmap of Jaccard similarities and (b) the resulting hierarchical clustering of the samples.

### 3.6 Runtime Performance

Finally we compare the runtime of the CONCOCT pipeline based on read mapping (which internally uses Bowtie2 [18], samtools [19], and Picard tools) to the GATTACA kmer counting implementation for estimating the coverage of pre-assembled contigs of a single HMP sample (SRS011134). The CONCOCT runtime is split as follows: 10.1 min for BWT index construction (from the contigs) and 51.2 min for coverage estimation in one sample. The GATTACA runtime consists of 5.3 min in kmer count index construction and 2.7 min for lookups in one sample. Therefore, ignoring the BWT indexing time, it would take CONCOCT roughly 81h to map the 95 HMP samples against these contigs; while GATTACA would require only 12.7h. However, if the samples in the cohort have already been indexed (for publicly available data or multi-sample studies reusing the same sample cohort), then GATTACA would only need 4.3h to finish (resulting in a roughly 20*×* speedup). The reported times were obtained running on a single 2.67 GHz Intel Xeon X5550 processor (in single-threaded mode). Both of the tools can be easily run in parallel across multiple samples, contigs, or sample reads.

## 4 Conclusion

GATTACA is a lightweight method for metagenomic binning, which achieves comparable binning accuracy with leading alignment-based methods at a fraction of the cost by approximating contig abundances using kmer counts. Furthermore, through offline indexing of publicly available cohorts of metagenomes, it enables efficient analysis of single metagenomic samples, catering to settings with scarce computational and data resources. As a result, an ungoing effort of this project is to index all the publicly-available samples from various metagenomic archives and make them available to the public.

